# Concise whole blood transcriptional signatures for incipient tuberculosis: A systematic review and patient-level pooled meta-analysis

**DOI:** 10.1101/668137

**Authors:** Rishi K. Gupta, Carolin T. Turner, Cristina Venturini, Hanif Esmail, Molebogeng X. Rangaka, Andrew Copas, Marc Lipman, Ibrahim Abubakar, Mahdad Noursadeghi

## Abstract

Blood transcriptional signatures may predict risk of tuberculosis (TB). We compared the performance of 17 mRNA signatures in a pooled dataset comprising 1,026 samples, including 183 samples from 127 incipient TB cases, from four studies conducted in South Africa, Ethiopia, The Gambia and the UK. We show that eight signatures (comprising 1-25 transcripts) that predominantly reflect interferon inducible gene expression, have equivalent diagnostic accuracy for incipient TB over a two-year period with areas under the receiver operating characteristic curves ranging from 0.70 (95% confidence interval 0.64-0.76) to 0.77 (0.71-0.82). The sensitivity of all eight signatures declined with increasing disease-free time interval. Using a threshold derived from two standard deviations above the mean of uninfected controls giving specificities of >90%, the eight signatures achieved sensitivities ranging 24.7-39.9% over a 24 month interval, rising to 47.1-81.0% over 3 months. Based on pre-test probability of 2%, the eight signatures achieved positive predictive value ranging from 6.8-9.4% over 24 months, rising to 11.1-14.3% over 3 months. When using biomarker thresholds maximising sensitivity and specificity with equal weighting to both, no signature met the minimum World Health Organization (WHO) Target Product Profile parameters for incipient TB biomarkers over a two-year period. Blood transcriptional biomarkers reflect short-term risk of TB and only exceed WHO benchmarks if applied to 3-6 month intervals.

## Introduction

Identification of people at high risk of developing tuberculosis (TB) enables the delivery of preventative treatment for a disease that accounts for more deaths than any other infectious disease worldwide, with 10 million incident cases and 1.6 million deaths estimated in 2017^1^. This approach represents a fundamental component of the World Health Organization (WHO) End TB strategy, aiming for a 95% reduction in TB mortality and 90% reduction in TB incidence by 2035^2^. However, these efforts are undermined by the poor predictive value (PPV) of current prognostic tests for development of TB, which focus on the identification of a T cell mediated response to mycobacterial antigen stimulation, as a surrogate for latent TB infection (LTBI)^3,4^. Poor prognostication of current diagnostics precludes precise delivery of preventative therapy, thus increasing costs and potential adverse effects, and attenuating the effectiveness of prevention programmes, while also limiting roll-out of preventative treatment to the limited-resource settings where the majority of TB cases occur.

Increasing recognition of the continuum of TB infection and disease has led to renewed interest in the ‘incipient’ phase of TB^5–7^. This has been defined by the WHO as the prolonged asymptomatic phase of early disease during which pathology evolves, prior to clinical presentation as active disease^8^. This definition encompasses the incipient and subclinical phases described by others^9^. This phase in the natural history of TB, between latent infection and active disease, is highly attractive as a target for novel prognostic TB diagnostic tests in the hope of improving PPV for incident TB, while still offering an opportunity to prevent TB-related morbidity and mortality and interrupt onward transmission^9^. This has led to the WHO producing a target product profile (TPP) for incipient TB diagnostics, stipulating minimum sensitivity and specificity of 75% and optimal sensitivity and specificity of 90%, over a two-year time period^8^. These minimum criteria are based on achieving PPV of 5.8%, when assuming 2% pre-test probability.

Multiple studies have discovered changes in the host transcriptome in association with TB disease, compared to healthy controls, individuals with LTBI or other diseases^10–15^. Signatures have become increasingly concise over time, making their translation to near-patient diagnostic tests more achievable. More recently, perturbation in the transcriptome has been found to predate the diagnosis of TB^14,16–18^, suggesting that transcriptional signatures may offer an opportunity to diagnose incipient TB and potentially fulfil the WHO TPP. However, independent validation of each signature is still limited. It remains unclear which of the multiple candidate transcriptional signatures performs best for the identification of incipient TB, or whether any signatures meet the WHO diagnostic accuracy benchmarks.

To address these critical knowledge gaps, we performed a comprehensive literature search to identify concise whole blood transcriptional signatures for incipient TB, along with whole blood transcriptomic datasets, with sampling prior to TB diagnosis. We then performed a patient-level pooled analysis to compare the diagnostic accuracy of the identified candidate transcriptional signatures for diagnosis of incipient TB among people at risk of disease over a two-year horizon. Finally, we evaluated the diagnostic accuracy of the best performing transcriptional signatures, stratified by pre-defined time intervals to TB, in order to critically assess their potential value as biomarkers in practice.

## Methods

### Systematic review

We performed a systematic review, in accordance with PRISMA standards^19^, to identify candidate concise whole blood transcriptional signatures for incipient or active TB, along with published whole blood transcriptomic datasets, with sampling prior to TB diagnosis. We searched Medline and Embase on 15/04/2019, with no language or date restrictions, using comprehensive MeSH and keyword terms (Supplementary Methods), and hand-searched reference lists of review articles. The pre-specified protocol for the review and analysis plan is available (PROSPERO CRD42019135618).

### Eligibility criteria for candidate signatures

We included whole blood mRNA signatures discovered with a primary objective of diagnosis of active or incipient TB, compared to asymptomatic controls. We included only signatures that used a defined approach to feature selection within genome wide data to reduce multidimensionality and lead to ‘concise’ signatures that may be more amenable to clinical translation. The availability of gene names that comprise the signature, along with the corresponding equation or modelling approach was required. We also specified that the signature (including component genes, and modelling approach) was validated in at least one independent test or validation set, in order to enable reliable signature reconstruction. We only included signatures discovered from training sets that included controls who were either deemed healthy, or had latent TB infection, since discriminating incipient TB from healthy or latently infected people is the primary aim of incipient TB diagnostics. Where multiple signatures were discovered for the same intended purpose and from the same training dataset, we included the signature with greatest accuracy, as defined by the area under the receiver operating characteristic curve (AUROC) in the validation data. Where accuracy was equivalent, we included the most parsimonious signature.

### Eligibility criteria for transcriptomic datasets

We included published whole blood transcriptomic datasets (RNAseq or microarray) where sampling prior to TB diagnosis was performed and interval time to disease was available. We specified a minimum median duration of follow-up of one year to reduce the risk of outcome misclassification. For studies where preventative TB therapy was offered, individual level data was required to identify the treated cases.

### Screening and data extraction

Two independent reviewers (RKG and CTT) screened titles and abstracts identified in the search, and determined eligibility for final inclusion following full-text review. Gene lists and corresponding equations or modelling approaches were extracted for each eligible candidate signature and validated by a second reviewer. Disagreements regarding study inclusion or signature calculations were resolved by a third reviewer (MN). Quality assessment and risk of bias were assessed for the studies corresponding to included RNA datasets, using modified versions of the Newcastle-Ottowa scale (using the cohort or case-control version as appropriate to each contributing study)^20^.

### Extension of UK cohort of TB case contacts

In preparation for this meta-analysis, we extended the follow-up of a previously published cohort of London TB contacts^18^, by relinking the full cohort to national TB surveillance records (until 31/12/2017) held at Public Health England, which includes all statutory national TB notifications (median follow up increased from 0.9 to 1.9 years). An additional 27 samples and individuals were also available for inclusion in the present analysis. The full updated data set for this study is available in ArrayExpress (Accession number awaited).

### RNA data processing

Individual level RNAseq data was downloaded for eligible studies, and mapped to the reference transcriptome (Ensembl Human GRCh38 release 95) using Kallisto^21^. The transcript-level output counts and transcripts per million (TPM) values were summed on gene level and annotated with gene symbols using tximport and BioMart^22,23^. Only protein-coding genes were selected for downstream processing, and TPM and counts per million (CPM) values <0.001 were set to 0.001 prior to log2 transformation. TPM data were compared for participants from different studies using principal component analysis (PCA) to test for heterogeneity and determine the need for batch correction. This included (a) the entire transcriptome; (b) selected genes comprising only the candidate signatures included in the analysis; and (c) invariant genes that were in the lowest quartile of genes ranked by variance within each of the contributing datasets. Batch correction was performed using the COmbat CO-Normalization Using conTrols (COCONUT) package in R^24^. This approach facilitated correction based on the disease-free controls, which was then applied to those with disease, thus reducing risk of bias during correction due to differing prevalence of disease among the study populations included.

### Definitions and sample inclusion

Only samples obtained prior to the diagnosis of TB were included. ‘Prevalent’ TB was defined as a TB diagnosis within 21 days of sample collection, as previously^4^. ‘Incipient TB’ cases were defined as individuals diagnosed with TB >21 days from blood RNA sample collection. Culture-confirmed and clinically or radiologically diagnosed pulmonary or extrapulmonary TB cases were included in the main analysis. ‘Non-progressors’ were defined as those who remain TB disease free during follow-up. Non-progressor samples with less than 6 months’ follow-up from the date of sample collection were excluded due to risk of outcome misclassification. Participants with prevalent TB and those who received preventative therapy were excluded. For studies with serial samples from the same individuals, serial samples were included provided that they met these criteria, and that they were collected at least 6 months apart, since they were treated as independent samples in the primary analysis.

### Calculation of signature scores

Scores were calculated for candidate signatures (using the authors’ described methods) for each participant in the pooled dataset. For signatures that required model reconstruction, we validated the reconstructed model against the original authors’ model, by comparing receiver operating characteristic (ROC) curves in their original test dataset when possible. Using a pre-defined ‘control’ population (including only participants with negative LTBI tests among the pooled dataset), batch-corrected signature scores were transformed to Z scores (by subtracting the control mean, and dividing by standard deviation), in order to standardise scaling across signatures^18^.

### Statistical analysis

ROC curves for each signature for the identification of incipient TB were first plotted, for a two-year time horizon. Any data that was originally used to derive specific signatures were excluded from the pooled dataset used to test the performance of the relevant signature. ROC curves and AUROCs for separate study datasets were initially examined visually to assess the between study heterogeneity. Since little heterogeneity was observed for all signatures, a simple pooled data analysis was performed thereafter. AUROCs were directly compared in a pairwise approach, using the DeLong method^25^. The best performing signature available from all samples in the pooled dataset was used as the reference for comparison with all other signatures. Correlation between signature scores was assessed using Spearman rank correlation. Pairwise Jaccard similarity indices between signatures were calculated using lists of their constituent genes. Clustered co-correlation and Jaccard index matrices were generated in Morpheus^26^ using average Euclidean distance. Upstream analysis of transcriptional regulation was performed using Ingenuity Pathway Analysis (Qiagen, Venlo, The Netherlands) and visualized as network diagrams in Gephi v0.9.2, depicting all statistically overrepresented molecules predicted to be upstream >2 target genes.

ROC curves and AUROCs were then assessed for the best performing signatures, using pre-specified intervals to TB of <3 months; <6 months; <1 year; and <2 years from sample collection. Sensitivity and specificity for each of these time intervals were determined at pre-defined cut-offs for each signature, defined as a standardised score of two (Z2), representing the 97.7th percentile of the IGRA-negative control population assuming a Normal distribution, as in previous work^18^. These estimates were used to model the estimated predictive values for incident TB across a range of pre-test probabilities.

We performed four sensitivity analyses. First, we restricted inclusion of TB cases to those with microbiological confirmation. Second, we included only one blood RNA sample per participant by randomly sampling one blood sample per individual, in studies which included serial sampling. We also examined sensitivity and specificity for the best performing signatures using the maximal Youden Index^27^ to achieve the highest accuracy within each time interval. Finally, we recomputed the ROC curves using mutually exclusive time intervals to TB of 0-3, 3-6, 6-12 and 12-24 months, for each curve excluding participants who had developed TB in an earlier interval.

## Results

### Systematic review process and summaries of included datasets and signatures

A total of 643 unique articles were identified in the systematic review (Supplementary Figure 1). Four RNA datasets (Table 1) and 17 signatures (Table 2) met the criteria for inclusion.

**Table 1:** Characteristics of the datasets included in meta-analysis of candidate whole blood transcriptional signatures for incipient tuberculosis (TB).

| Study | Sample size | Study design | Population | Setting | HIV status | Sampling | Follow-up duration | TB case definition | RNAseq methods | NOS score | Baseline TB assessment |
| --- | --- | --- | --- | --- | --- | --- | --- | --- | --- | --- | --- |
| **London TB contacts**<sup>18</sup> | 360 | Cohort | Adult TB contacts | London | Negative | Baseline | 1.9 years (median) | Culture-confirmed, or clinically-diagnosed | 15-20 million 41 base pair paired-end reads | 7 (/7) | Clinical evaluation and chest radiograph |
| **Adolescent Cohort Study**<sup>16</sup> | 355 | Nested case-control | Adolescents with latent TB infection | South Africa | Negative | Serial (0, 6, 12, 24 months) | 2 years | Intrathoracic disease with 2 positive smears, or 1 positive culture | 30 million 50 base pair paired-end reads | 9 (/9) | Clinical evaluation. TB <6 months from enrolment excluded. Chest radiograph not specified |
| **Grand Challenges 6-74**<sup>17</sup> | 434 | Nested case-control | Adult household pulmonary TB contacts | South Africa, The Gambia, Ethiopia | Negative | Serial (0, 6, 18 months) | 2 years | Culture-confirmed, or clinically-diagnosed | 60 million 50 base pair paired-end reads | 9 (/9) | Clinical evaluation. TB <3 months from enrolment excluded. Chest radiograph not specified |
| **Leicester TB contacts**<sup>14</sup> | 108 | Cohort | Adult TB contacts | Leicester | Negative | Baseline + serial for a subset* | 2 years | Culture- or Xpert MTB/RIF-confirmed | 25 million 75 base pair paired-end reads | 7 (/7) | Clinical evaluation and chest radiograph |

*\*Due to the high frequency of serial sampling (<6-monthly), only baseline samples were included. NOS = Newcastle-Ottawa Scale (denominators shown in* *brackets).*

**Table 2.**
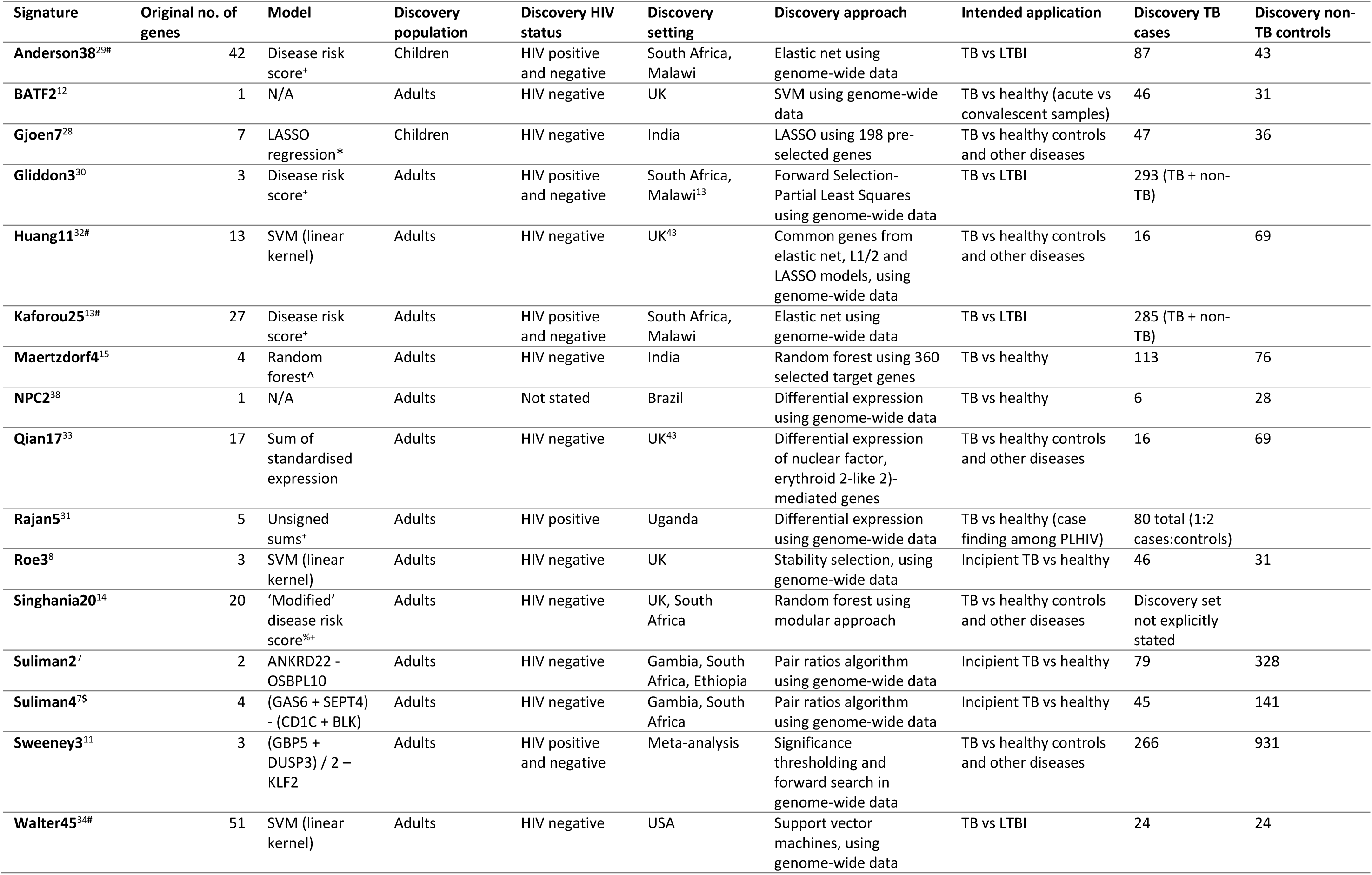
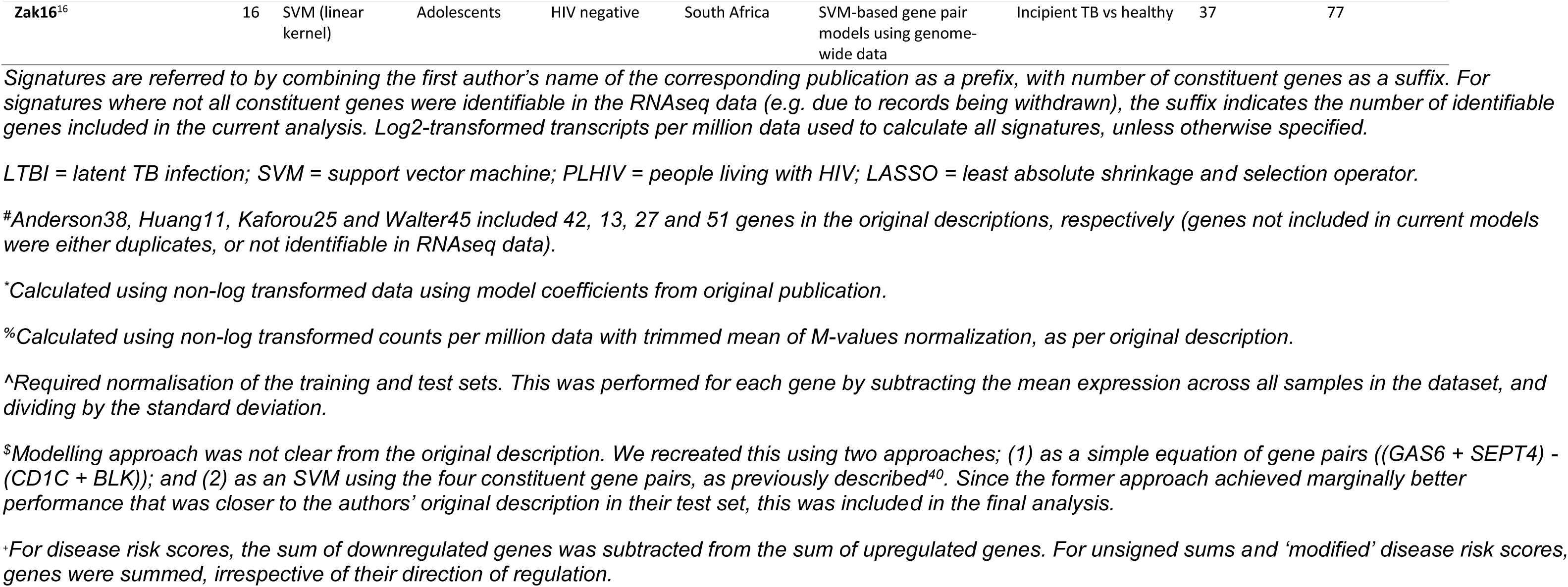
Characteristics of candidate whole blood transcriptional signatures for incipient tuberculosis (TB) included in systematic review and meta-analysis.

The RNA datasets included the Adolescent Cohort Study (ACS) of South African adolescents with LTBI^16^, the Bill and Melinda Gates Foundation Grand Challenges 6-74 (GC6-74) household TB contacts study in South Africa, the Gambia and Ethiopia^17^, a London TB contacts study^18^, and a Leicester TB contacts study^14^. The ACS and GC6-74 studies were nested case-control designs within larger prospective cohort studies, while the London and Leicester TB contacts studies were prospective cohort studies. All four studies were done in HIV-negative participants. The London TB contacts study included only baseline samples, while the ACS, GC6-74 and Leicester TB contacts studies included serial sampling. All four studies achieved maximal quality assessment scores.

A total of 1,126 samples from 905 patients met our criteria for inclusion (Supplementary Figure 2). These included 183 samples from 127 incipient TB cases, of which 117 (92.1%) were microbiologically confirmed. Baseline characteristics of the study participants are shown in Supplementary Table 1. Principal component analyses (PCA) revealed clear separation of samples by dataset when including (a) the entire transcriptome; (b) selected genes comprising only the candidate signatures included in the analysis; and (c) invariant genes, indicative of batch effects in the data that were eliminated after batch correction (Supplementary Figure 3).

Of the 17 identified signatures (Table 2), two were discovered from paediatric populations^28,29^. Four signature discovery datasets included HIV-infected and –uninfected participants^11,13,29,30^, one was discovered in an exclusively HIV-infected population for the purpose of active case finding^31^, while the remainder were discovered in HIV-negative populations. Four signatures were discovered with the intention of diagnosis of incipient TB^16–18^, with the remaining 13 discovered for diagnosis of active TB disease. Of these, five^11,14,28,32,33^ targeted discrimination of TB from other diseases in addition to discriminating people who were healthy or with LTBI. Of the 17 included signatures, only three were not discovered through a genome-wide approach^15,28,33^. Four signatures required reconstruction of support vector machine (SVM) models^16,18,32,34^, and one required reconstruction of a random forest model^15^. Our reconstructed models were validated against the authors’ original descriptions by comparing AUROCs in common datasets (Supplementary Table 2). The distribution of signature scores, stratified by study, pre- and post-COCONUT batch correction is shown in Supplementary Figure 4.

### Eight signatures perform equivalently for identification of incipient TB over two-year horizon

We first examined ROC curves and corresponding AUROCs for the identification of incipient TB by all 17 signatures over a two-year period in the separate contributing study datasets (Supplementary Table 3). This analysis initially suggested overall lower AUROCs in the GC6-74, compared to the ACS dataset. However, the distribution of TB events during follow-up differed between these studies (Supplementary Table 1). Following stratification by interval to disease, similar AUROCs were observed between studies, suggesting that interval to disease confounded the association between source study and AUROC. Since little residual between study heterogeneity was observed and PCA post-batch correction showed no clustering by study (Supplementary Figure 3), we proceeded to perform a pooled data analysis without further adjustment for source study.

We omitted scores for the Suliman2, Suliman4 and Zak16 signatures for their corresponding training sets within the GC6-74 and ACS datasets. The signature with highest AUROC for the identification of incipient TB over a two-year period tested in pooled data from all 1,126 samples was BATF2 (AUROC 0.74; 95% confidence interval 0.69-0.78). BATF2 was therefore used as the reference standard for paired comparisons of the other 16 candidate signatures. We found that eight signatures had equivalent AUROCs. These were Suliman2 (AUROC 0.77; 95% CI 0.71-0.82), BATF2 (0.74; 0.69-0.78), Kaforou25 (0.73; 0.69-0.78), Gliddon3 (0.73; 0.68-0.77), Sweeney3 (0.72; 0.68-0.77), Roe3 (0.72; 0.67-0.77), Zak16 (0.7; 0.64-0.76) and Suliman4 (0.7; 0.64-0.76) (Supplementary Table 4). The remaining nine signatures had significantly inferior AUROCs.

### The best performing RNA signatures are highly correlated

Next, we examined the correlation between the 17 candidate signature scores in the pooled dataset, as defined by Spearman rank correlation. The eight signatures identified with equivalent performance demonstrated moderate to high correlation (Supplementary Figure 5; correlation coefficients 0.44-0.84). In contrast, Singhania20, Anderson38, Huang11 and Walter45 showed little correlation with any other signature. The correlation matrix dendrogram showed closest relationships between signatures identified by the same research group.

To assess whether correlation was driven by overlapping constituent genes, we calculated pairwise Jaccard Indices (Supplementary Figure 5). There was a weak positive association between Spearman rank correlation and Jaccard Index, suggesting that overlapping constituent genes may partially account for their correlation. Genes comprising the eight signatures with equivalent AUROCs are demonstrated in Figure 1. Upstream analysis predicted that interferon-gamma, STAT1, interferon-alpha and tumour necrosis factor were the strongest predicted transcriptional regulators of these constituent genes.

**Figure 1.**
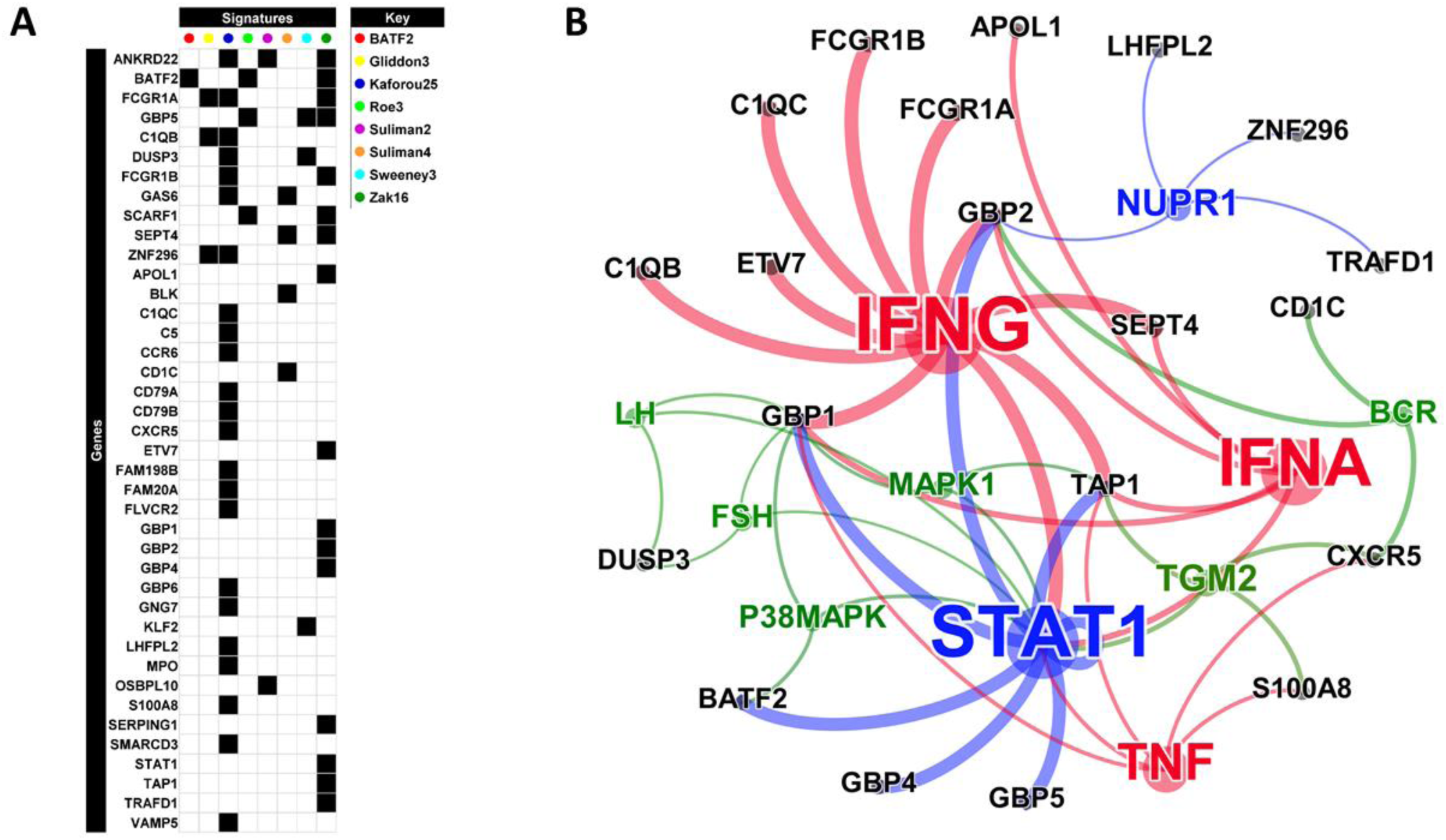
Genes comprising the top 8 blood transcriptomic signatures for incipient tuberculosis (TB) shown as (a) matrix; and (b) network diagram. Network diagram shows statistically enriched (p<0.05) upstream regulators of the 40 genes, identified by IPA. Coloured nodes represent the predicted upstream regulators, grouped by function (red= cytokine, blue=transcription factor, green=other). Grey nodes represent the transcriptional biomarkers downstream of these regulators. The identity of each node is indicated using Human Genome Organisation (HUGO) nomenclature. The size of the nodes is proportional to the number of downstream biomarkers associated with each regulator and the thickness of the edges is proportional to the - log10 P value for enrichment of each of the upstream regulators.

### Diagnostic accuracies of candidate signatures decline with interval to disease

Scores of the eight best performing signatures, stratified by interval to disease, are shown in Figure 2. AUROCs of these signatures declined with increasing interval to disease (Figure 3), ranging 0.82-0.91 for 0-3 months *vs.* 0.73-0.82 for 0-12 months.

**Figure 2.**
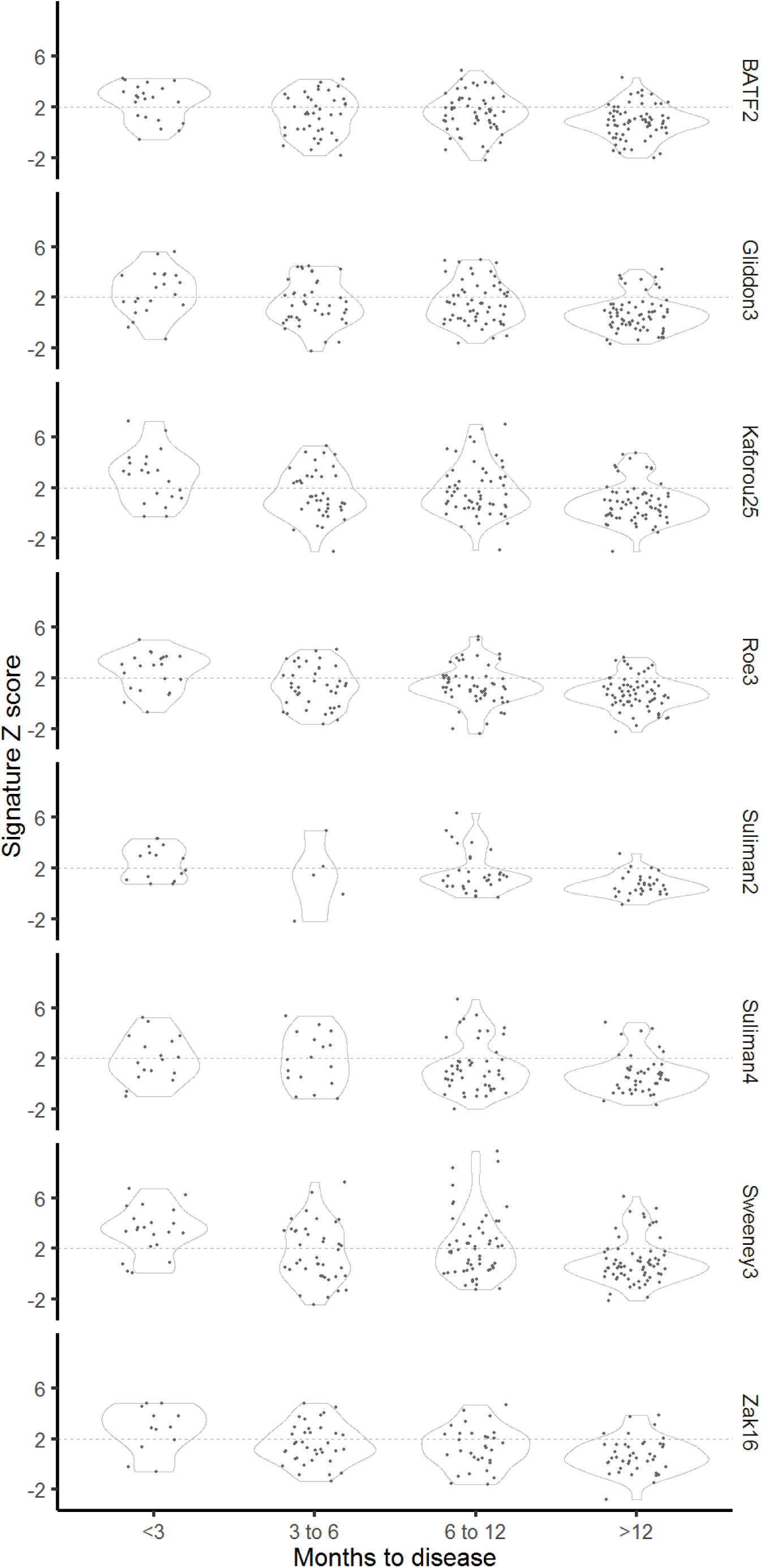
Scatterplots showing scores of eight best performing transcription signatures for incipient tuberculosis (TB), stratified by interval to disease. Dashed horizontal lines indicate thresholds set as standardised scores of two (Z2) for each signature.

**Figure 3.**
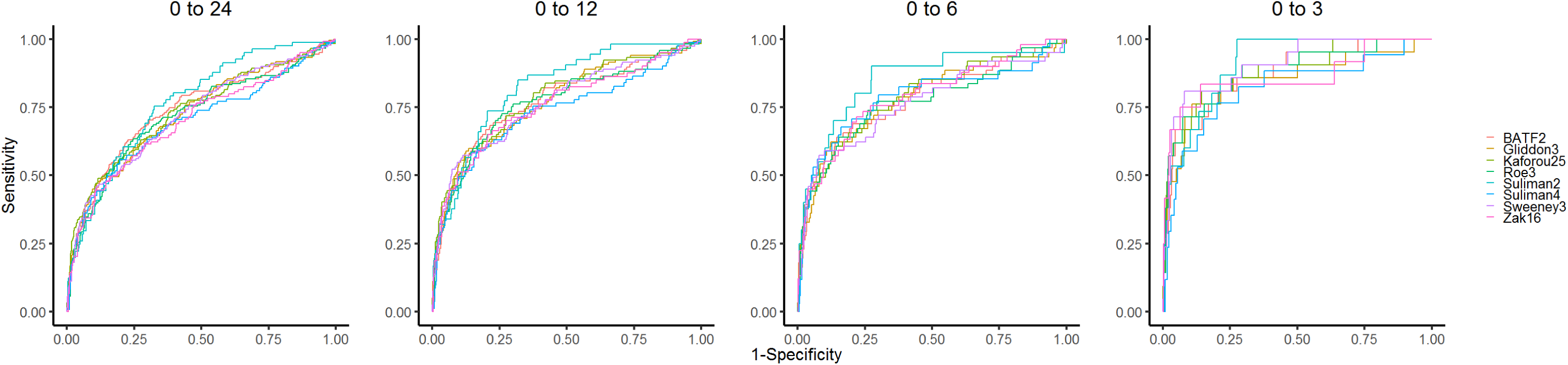
Receiver operating characteristic curves showing diagnostic accuracy of eight best performing transcriptional signatures for incipient tuberculosis (TB), stratified by months from sample collection to disease.

Figure 4 demonstrates diagnostic accuracy of the eight best performing candidates using pre-specified Z2 cut-offs based on the 97.7th percentile of the IGRA-negative control population, stratified by interval to disease and benchmarked against positive predictive value (PPV) estimates based on a pre-test probability of 2%. At this threshold, test sensitivities over 0-24 months of the eight best performing signatures ranged from 24.7% (16.6-35.1) to 39.9% (33.0-47.2) for the Suliman2 and Sweeney3 signatures, respectively, while corresponding specificities ranged from 92.3% (89.8-94.2) to 95.3% (92.3-96.9). In contrast, over a 0-3 month interval, sensitivities ranged from 47.1% (26.2-69.0) for the Suliman4 signature to 81.0% (60.0-92.3) for the Sweeney3 signature, with corresponding specificities of 90.9% (88.9-92.6) to 94.8% (93.0-96.2). For each of the time points, each of the eight signatures fell in the same PPV plane (5-10% over 0-24 months *vs.* 10-15% over 0-3 months).

**Figure 4.**
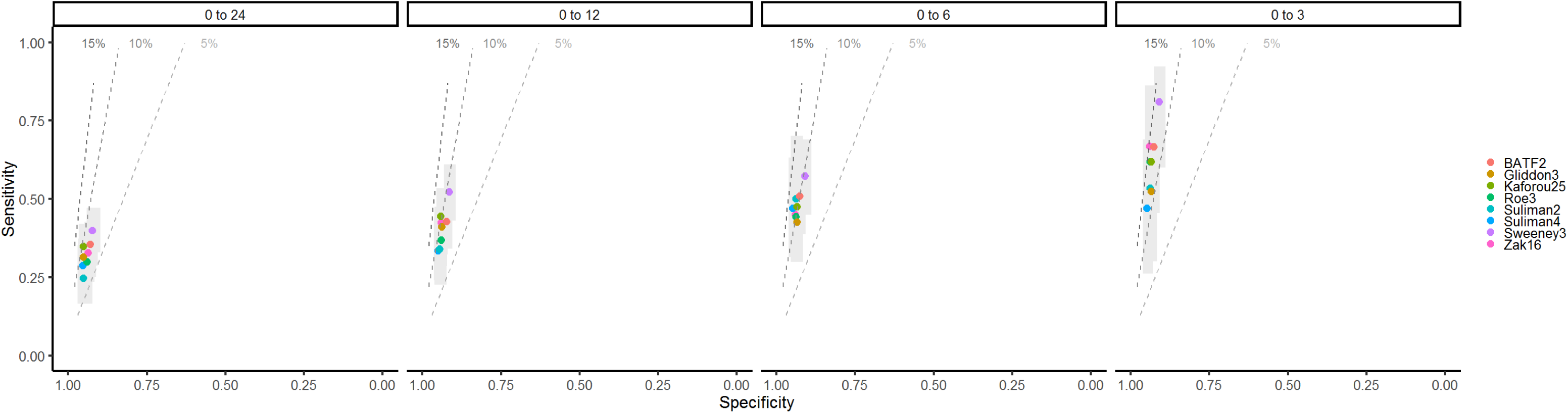
Diagnostic accuracy of eight best performing transcriptional signatures for incipient tuberculosis (TB) shown in receiver operating characteristic space, stratified by months to disease. Dashed lines represent positive predictive value planes, based on 2% pre-test probability. Grey shaded zones indicate 95% confidence intervals for each signature. Raw data presented in Supplementary Table 5.

### Predictive values for incipient TB

Positive- and negative-predictive values, modelled across a range of pre-test probabilities, are shown in Supplementary Figure 6. Based on pre-test probability of 2% all eight top performing signatures achieved a PPV marginally above the WHO benchmark of 5.8% for a 0-24 month period, ranging from 6.8% for Suliman2 to 9.4% for Kaforou25, with corresponding NPVs of 98.4% and 98.6%, respectively. For the 0-3 month time period, PPVs ranged from 11.1% for Gliddon3 to 14.3% for Zak16, with corresponding NPVs of 99.0% and 99.3%, respectively.

### Sensitivity analyses

Restricting inclusion of incipient TB cases to those with documented microbiological confirmation and including only one blood RNA sample per participant (by randomly sampling) produced no significant change to the main results (Supplementary Figures 7 and 8). Sensitivities and specificities of the top eight equivalent signatures using cut-offs defined by the maximal Youden index for each time interval fell below the minimum WHO TPP criteria for a 0-24 month period, but met or approximated the minimum criteria over 0-3 months (Supplementary Figure 9). Finally, reanalysis of the ROC curves using mutually exclusive time periods of 0-3, 3-6, 6-12 and 12-24 months magnified the difference in performance between the intervals, with performance declining more markedly with increasing interval to disease (Supplementary Figure 10).

## Discussion

In the largest analysis to date of the performance of whole blood transcriptional signatures for incipient TB, we demonstrate that eight candidate transcriptional signatures performed with equivalent diagnostic accuracy for incipient TB over a two-year period. These signatures ranged from a single transcript (BATF2) to 25 genes (Kaforou25). The accuracy of all eight signatures declined markedly with increasing intervals to disease, suggesting that they represent biomarkers of short-term risk of disease. When benchmarked against the WHO TPP for incipient TB biomarkers over a two-year period, these signatures only marginally surpassed the target PPV of 5.8%, when assuming 2% pre-test probability and using a cut-off of two standard scores (Z2). However, sensitivity at this Z2 cut-off was only 24.7-39.9%, with the majority of cases being missed. Moreover, no signature achieved the target sensitivity and specificity of ≥75% over two years, even when using the cut-off with maximal accuracy (as defined by the Youden Index) for this time period for each signature individually. In contrast, the eight best performing signatures achieved PPVs of 11.1-14.3%, NPVs >98.9% and sensitivities of 47.1-81.0% over a 0-3 month period, assuming the same pre-test probability, suggesting that exceeding performance of the WHO benchmarks is likely more achievable over a shorter time frame.

### Eight signatures have equivalent diagnostic accuracy for incipient TB

In light of increasing industry interest in translation to clinically implementable platforms, our findings have important commercial implications. The cost of transcriptional signatures is highly unlikely to match the minimal target price of <$2 specified by WHO TPP for a non-sputum triage test for TB disease^35^. The most likely application of this technology is therefore likely to be diagnosis of incipient TB, where the WHO TPP pricing parameters are more lenient, aiming for an initial price of <$100, with the price of IGRAs as an initial benchmark^8^. The application of these biomarkers for incipient TB diagnosis also mitigates the major limitation of the imperfect specificity of predominantly interferon-inducible gene signatures when trying to discriminate TB from other infectious diseases, by applying them to a predominantly healthy and asymptomatic population^36^. The equivalence of diagnostic accuracy of eight signatures for incipient TB in this analysis suggests that parsimony of genes, stability of cross-platform transcript measurements, and simplicity of modelling calculations may be key to informing selection of signatures from these candidates for further validation.

The eight signatures that achieved equivalent performance were discovered with the primary intention of diagnosis of incipient TB^16–18^, or differentiating active TB from people who are healthy or with LTBI ^12,13,30,37^. Discovery populations for these eight signatures included adults or adolescents from the UK or Southern Africa^12,13,16–18,30^, or a meta-analysis of microarray data from multiple studies^11^, including a minimum of 37 incipient or active TB cases. All eight signatures were discovered using genome-wide approaches. Correlation analysis revealed moderate to high correlation between all eight of these signatures, while the 40 genes comprising these signatures were predominantly driven by interferon-gamma, STAT1, interferon-alpha and TNF. This evidence suggests that these signatures reflect a common host immune response, which explains their equivalent performance. Signatures identified by the same research group clustered together within the correlation dendrogram, which likely reflects common modelling approaches and study populations used during discovery within the same research group.

In contrast, the nine signatures with inferior performance included two discovered from children^28,29^, one study that prioritised discrimination of active TB from other bacterial and viral infections^14^, and one study that conducted active case-finding for TB among people living with HIV^31^. The differences in primary intended applications, which are reflected in the study populations used for biomarker discovery, may account for their inferior performance when evaluated solely for identification of incipient TB in a predominantly healthy, HIV-negative adult population. The signatures with inferior performance also included three discovered from panels of pre-selected candidate genes, rather than a genome-wide approach^15,28,33^, and four with only 6-24 TB cases in the discovery sets^32–34,38^. This suggests that using a genome-wide approach and including adequate numbers of diseased cases are important considerations during signature discovery.

### Diagnostic accuracy is time-dependent

The time-dependent performance of the signatures is likely explained by their expression being a reflection of an underlying disease process of incipiency. If true, this suggests that the duration of the incipient phase of TB is typically less than three months. Lower sensitivity at longer intervals to disease may be explained by commencement of the incipient phase and/or re-exposure to TB between sampling and diagnosis. However, even for a 0-3 month interval, sensitivity ranged from 47.1-81.0% for the eight best performing signatures, with some cases missed. This observation may be explained by truly imperfect signature sensitivity for the incipient phase, or very rapid disease progression among a subset of cases.

In order to achieve the WHO TPP, a screening strategy that incorporates serial testing on a 3-6 monthly basis may therefore be required for transcriptional signatures. Such a strategy, however, is unlikely to be feasible at a population level. Instead, high-risk groups such as household contacts could be targeted, though even this is still unlikely to be scalable in high transmission settings, given the limited global coverage of contact-tracing programmes. In lower transmission, higher resource settings, serial blood transcriptional testing for risk-stratification over a defined 1-2 year period may be more achievable, particularly among recent contacts or new entry migrants from high transmission countries, for whom risk of disease is highest within an initial two-year interval^4,18,39^.

### Strengths and weaknesses

Strengths of this study include the size of the pooled dataset, including 1,126 samples from 905 patients, and 183 samples from 127 incipient TB cases. All four contributing studies achieved maximal quality assessment scores and were performed in relevant target populations of either recent TB contacts, or people with LTBI. This facilitated a robust analysis of diagnostic accuracy of the candidate signatures, stratified by interval to disease. Second, we performed a comprehensive systematic review, and identified 17 candidate signatures. For each of these signatures, gene lists and modelling approaches were extracted and validated by independent reviewers. Moreover, for signatures that required model reconstruction, our models were cross-validated against original models by comparing AUROCs using the same dataset wherever possible. This allowed us to perform a comprehensive, head-to-head analysis of candidate signatures for incipient TB for the first time, ensuring that each head-to-head comparison was performed on paired data. This contrasts with a recent head-to-head systematic evaluation that included only two of the eight best-performing signatures in our analysis, and compared performance for incipient TB in only one dataset over a 0-6 month time period^40^. Finally, our meta-analytic methods ensured a standardised approach to RNAseq data. This included an unbiased approach to batch correction, with unchanged distributions of signature scores within each dataset following correction.

A weakness of our analysis is that we were unable to perform subgroup analyses by age, ethnicity or country, since the contributing studies largely defined these strata. Reassuringly, there were no clear differences in performance by study, supporting the generalisability of the results. Secondly, having observed little heterogeneity between studies, we conducted a simple pooled analysis. The precision of our estimates therefore may be slightly overstated and statistical tests may be anti-conservative. Likewise, treating serial samples as independent was anti-conservative, but findings were similar in our sensitivity analysis taking only one sample per individual at random.

All included datasets were from the UK or sub-Saharan Africa, while no data were available for people living with HIV or children under 10, among whom different blood transcriptional perturbations may occur in TB^5,29^. Prospective validation studies in other world regions and among these specific target populations are needed. In addition, the majority of incipient TB cases were contributed from the African datasets, with 12 cases from the UK studies. Nevertheless, the UK studies were done in appropriate target populations of close contacts of TB index cases and were performed as cohort studies, as opposed to the African case-control designs. High specificity for correctly identifying non-progressors among contacts is a key attribute in improving PPV, compared to existing tests. Hence, these UK datasets were extremely valuable additions to the pooled meta-analysis.

Finally, while every effort was made to reconstruct signature models according to authors’ descriptions, small differences in our reconstructions may have led to under- or over-estimation of performance. However, validation of our reconstructed models against the authors’ original models showed reassuringly minimal difference in AUROCs using common datasets, suggesting the impact of this bias is likely very small.

### Future directions

Future studies addressing transcriptional biomarkers for incipient TB may address diagnostic accuracy and predictive value, or the impact of these biomarkers on health system and patient outcomes, as previously outlined^41^. Studies further assessing accuracy and predictive value should continue to be performed among relevant target populations, with sufficient follow-up for disease progression, and should ensure that TB diagnosis is ascertained blind to biomarker results. These may include further head-to-head analyses of candidate signatures, particularly addressing population subgroups and world regions beyond the UK and sub-Saharan Africa. In contrast, prospective impact studies may incorporate delivery of preventative treatment based on transcriptional biomarker results to further assess the feasibility, effectiveness and cost-effectiveness of transcriptional biomarker-stratified preventative therapy. Approaches may include a two-step algorithm, where serial testing is only performed among patients with LTBI, in order to rationalise the burden and costs of follow-up. Such prospective studies, with real time delivery of biomarker results, would also allow prospective clarification of the clinical significance of transcriptional signature expression, by ensuring the absence of clinical, radiological or microbiological evidence of TB disease among participants with a positive test. However, clinical evaluations to assess for such evidence should not surpass those performed under routine programmatic conditions^41^. Moreover, integral to both diagnostic accuracy and impact studies is the translation of transcriptional measurements from un-scalable genome-wide approaches such RNAseq to the reproducible quantification of selected signature genes, with appropriately defined cut-offs. While this has been performed for some signatures using PCR-based platforms^16,17,30,42^, no signature platforms have yet been validated for implementation in a near-patient or commercial assay.

## Conclusions

In summary, we demonstrate for the first time that eight transcriptional signatures, including a single transcript (BATF2), have equivalent diagnostic accuracy for identification of incipient TB. Performance appeared similar across studies, including participants from the UK and Southern Africa. Signature performance was highly time-dependent, with lower accuracy at longer intervals to disease. A screening strategy that incorporates serial testing on a 3-6 monthly basis among selected high-risk groups may be required for these biomarkers to surpass WHO TPP benchmarks.

## Author Contributions

RKG, IA and MN conceived the study. RKG, CTT and MN wrote the systematic review protocol, performed the literature review and extracted signature models. RKG performed the analyses and wrote the first draft of the manuscript, supported by CTT and MN. All other authors contributed to the methods or interpretation. All authors have seen and agreed on the final submitted version of the manuscript.

