## Supplemental appendix for "Concise whole blood transcriptional signatures for incipient tuberculosis: A systematic review and patient-level pooled meta-analysis"

### Supplementary Appendix

**Supplementary Methods: Details of Medline search strategy**

1. Biomarkers/
2. Diagnostic Tests, Routine/
3. "Predictive Value of Tests"/
4. diagnostic test*.mp.
5. biomarker*.mp.
6. ppv.mp.
7. npv.mp.
8. sensitivit*.mp.
9. specificit*.mp.
10. signature*.mp.
11. 1 or 2 or 3 or 4 or 5 or 6 or 7 or 8 or 9 or 10
12. exp TUBERCULOSIS/ or tuberculosis.mp. or exp MYCOBACTERIUM TUBERCULOSIS/ or tb.mp.
13. RNA/
14. Transcriptome/
15. rna.mp.
16. transcript*.mp.
17. gene expression.mp.
18. Gene Expression Profiling/ or RNA, Messenger/ or Transcription, Genetic/ or Gene Expression/
19. 13 or 14 or 15 or 16 or 17 or 18
20. 11 and 12 and 19
21. blood/
22. blood.mp.
23. 21 or 22
24. 11 and 12 and 19 and 23
25. remove duplicates from 24
26. limit 25 to "humans only (removes records about animals)"

**Supplementary Figure 1.** Flowchart showing systematic review process for review and meta-analysis of concise whole blood transcriptional signatures for incipient tuberculosis.

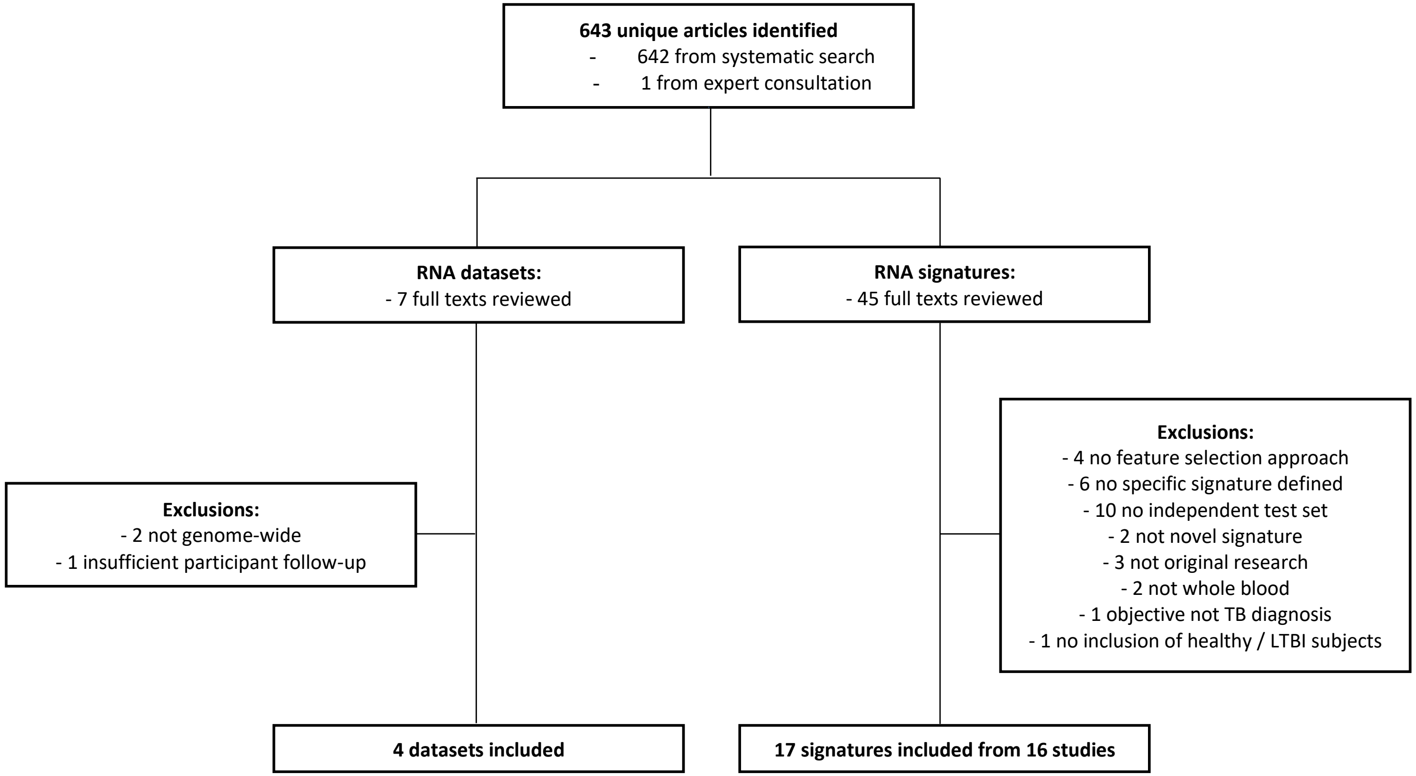

**Supplementary Figure 2.** Inclusion of samples from contributing datasets in meta-analysis of concise whole blood transcriptional signatures for incipient tuberculosis (TB). ACS = adolescent cohort study; GC6-74 = Bill and Melinda Gates Foundation Grand Challenges 6-74 TB contacts study; PT = preventative therapy.

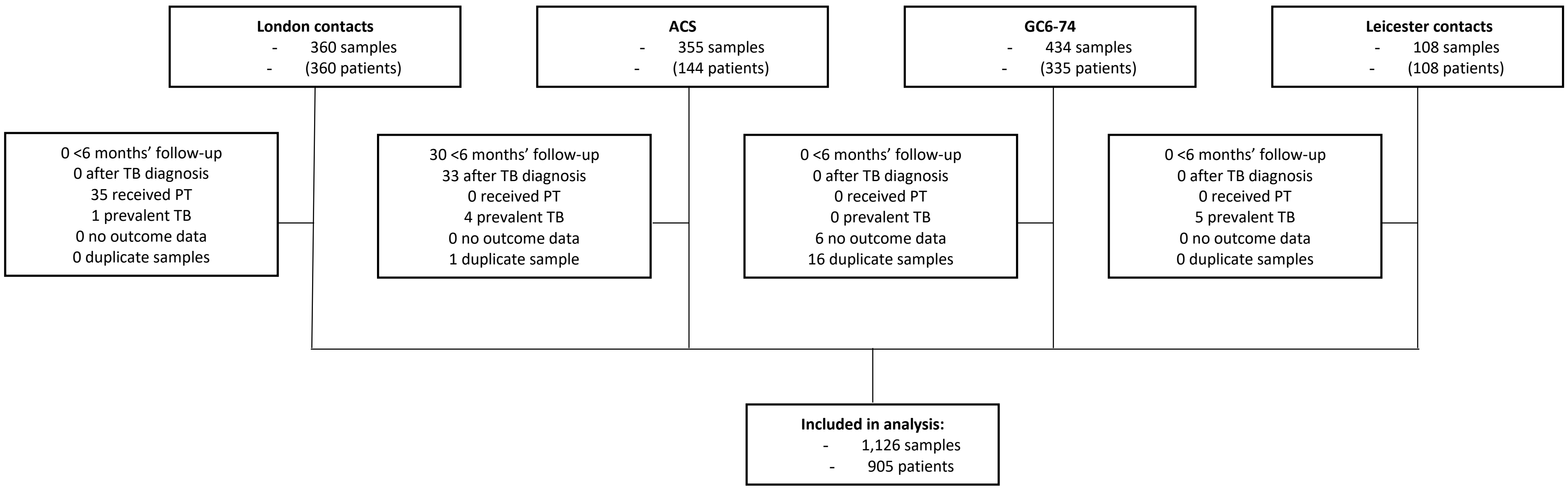

**Supplementary Figure 3.** Principal component analysis (PCA) of RNAseq before batch correction showing (a) the entire transcriptome; (b) selected genes comprising only the candidate signatures included in the analysis; and (c) invariant genes, stratified by source study. Panel (d) shows PCA following batch correction. ACS = adolescent cohort study; GC6-74 = Bill and Melinda Gates Foundation Grand Challenges 6-74 TB contacts study.

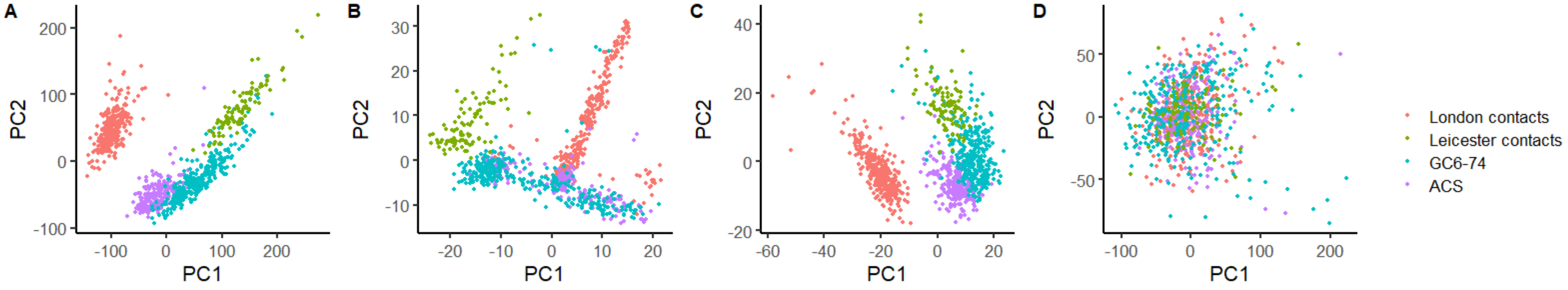

**Supplementary Figure 4.** Density plots of signature expression (a) before and (b) after batch correction, stratified by source study. ACS = adolescent cohort study; GC6-74 = Bill and Melinda Gates Foundation Grand Challenges 6-74 TB contacts study.

**A**

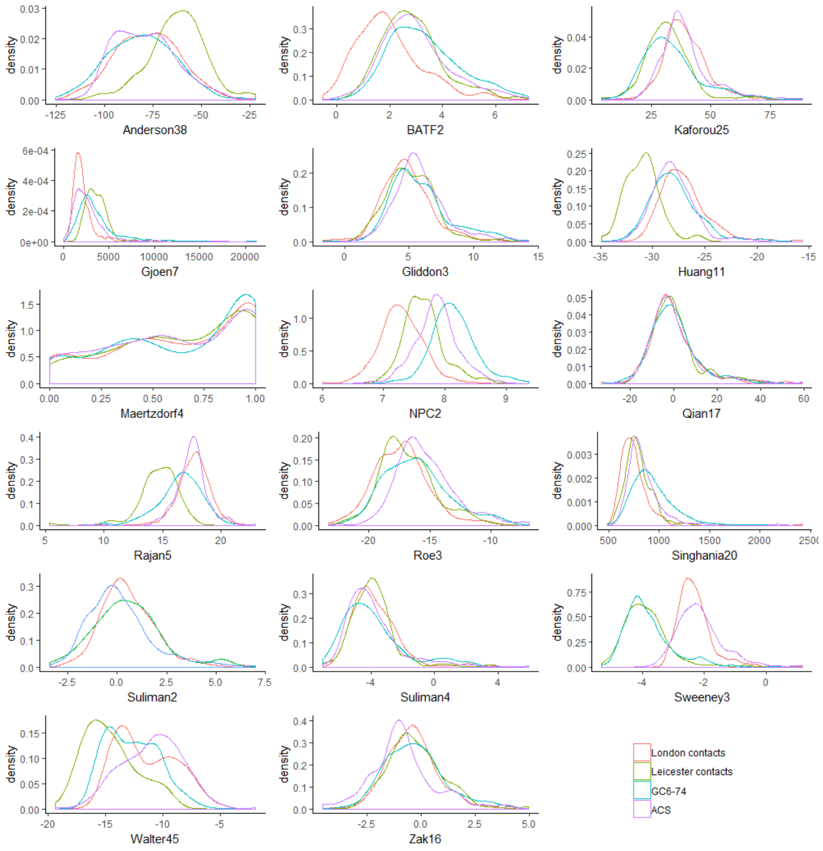


**B**

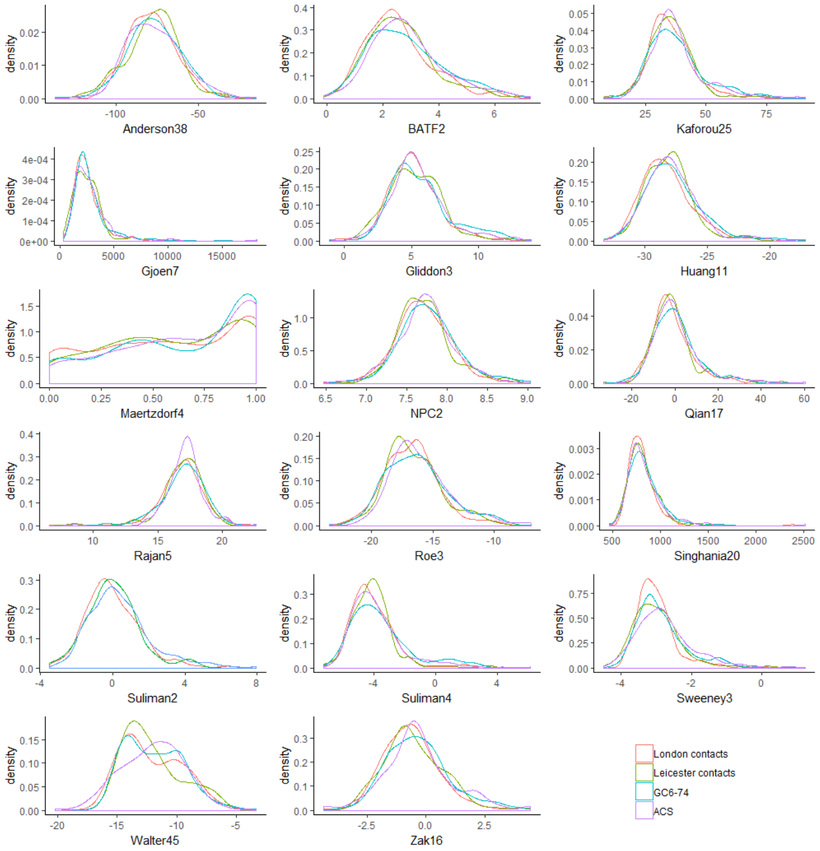

**Supplementary Figure 5.** Relationships between 17 candidate transcriptional signatures for incipient tuberculosis (TB) included in meta-analysis displayed as: (a) Spearman rank correlation matrix heatmap; (b) Jaccard similarity index heatmap showing overlapping constituent genes; and (c) Jaccard index vs. Spearman correlation coefficient for pairwise signature comparisons.

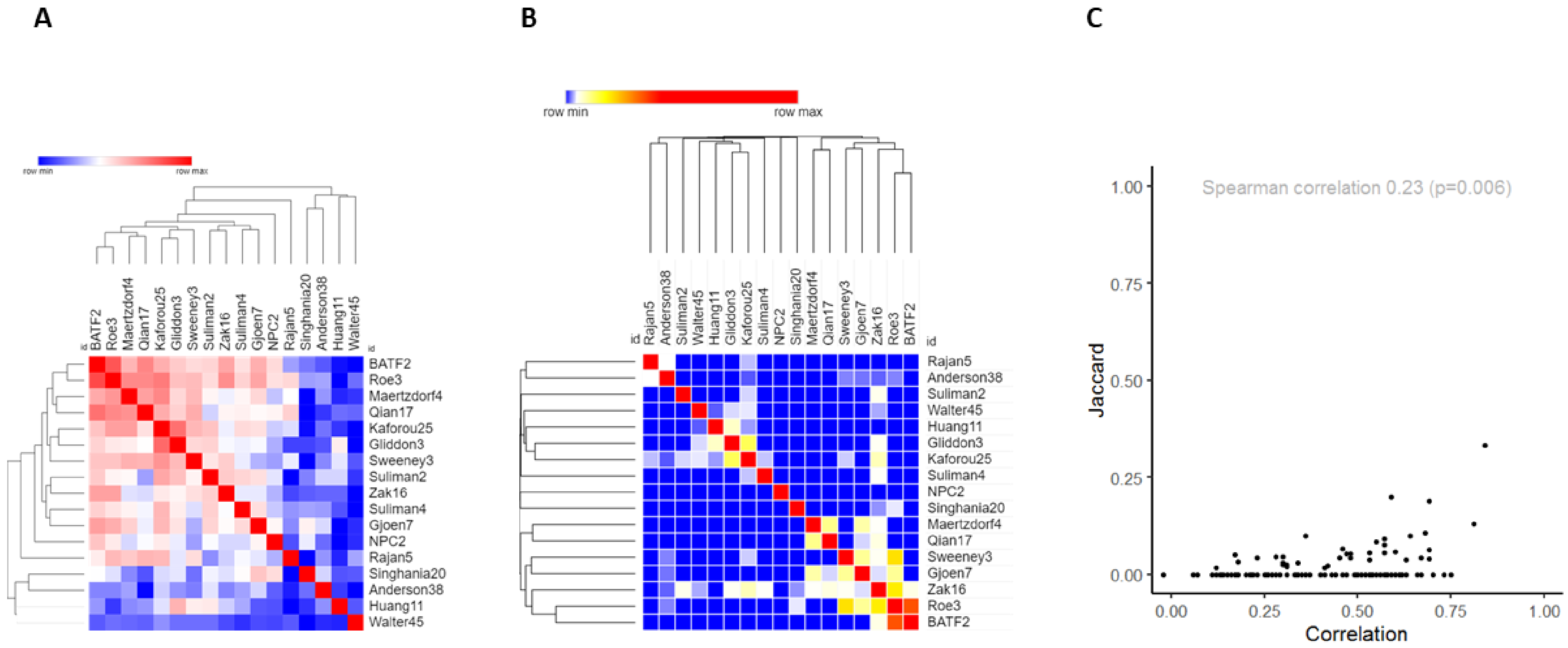

**Supplementary Figure 6.** Positive- and negative-predictive values (PPVs/NPVs), modelled across a range of pre-test probabilities for eight best performing transcriptional signatures for incipient tuberculosis (TB), and stratified by months to disease. Dashed line indicates 2% pre-test probability.

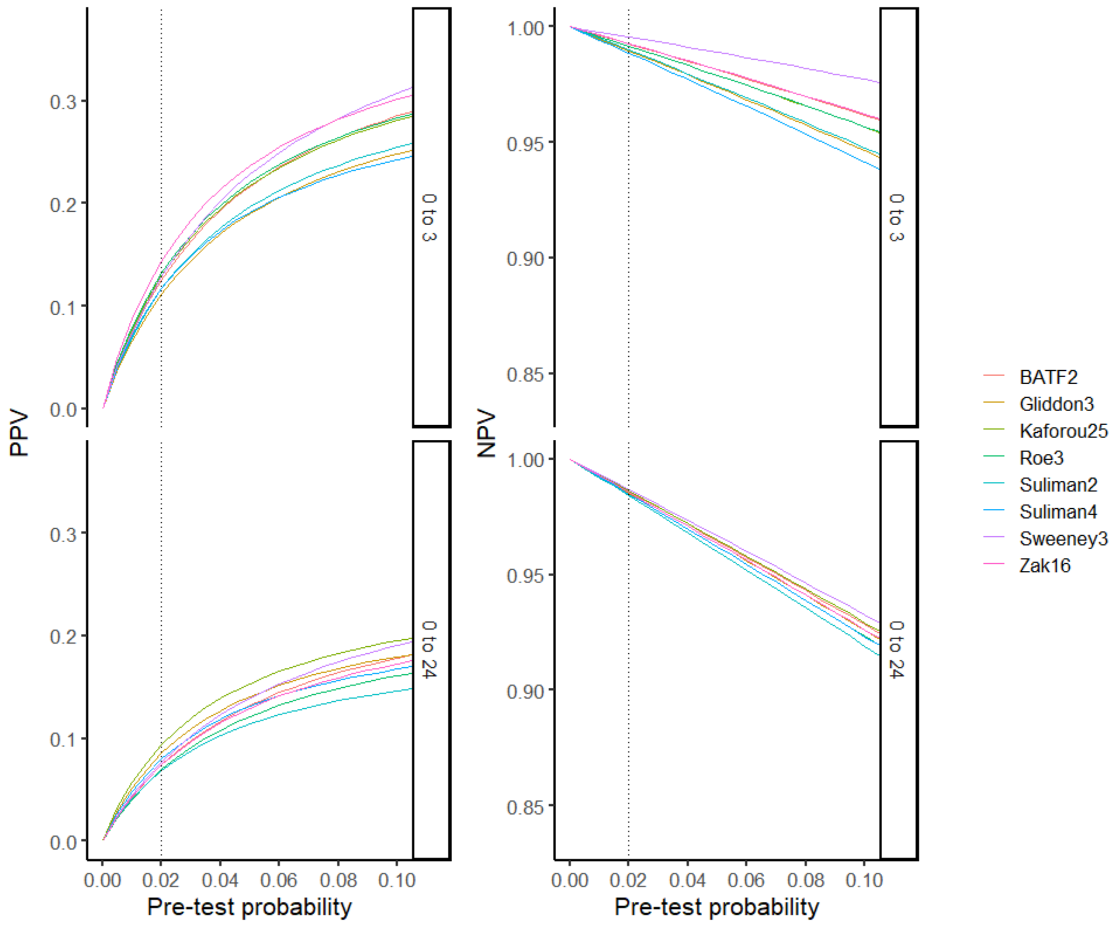

**Supplementary Figure 7.** Receiver operating characteristic curves showing diagnostic accuracy of eight best performing transcriptional signatures for incipient tuberculosis (TB), stratified by months from sample collection to disease. Sensitivity analysis restricting inclusion of incipient TB cases to those with documented microbiological confirmation.

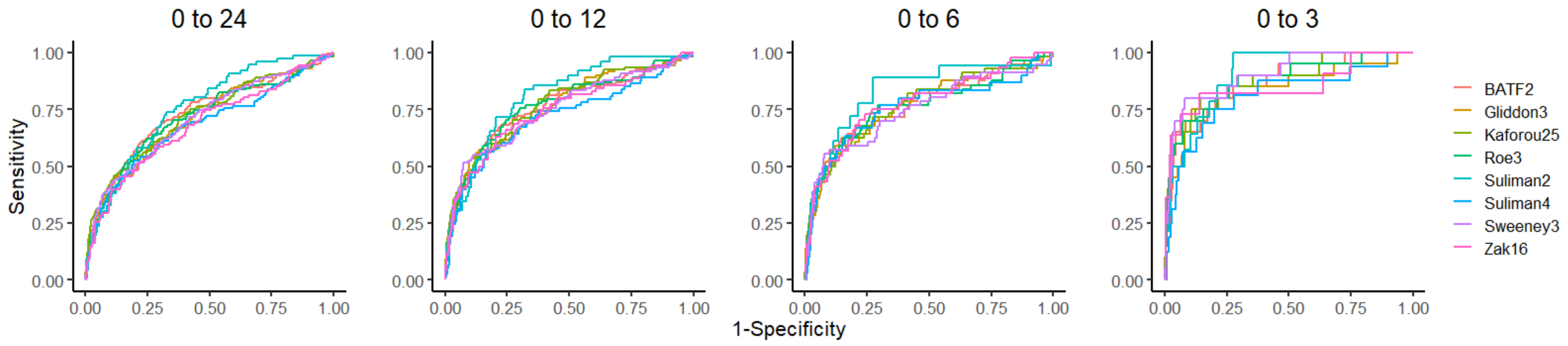

**Supplementary Figure 8.** Receiver operating characteristic curves showing diagnostic accuracy of eight best performing transcriptional signatures for incipient tuberculosis (TB), stratified by months from sample collection to disease. Sensitivity analysis including only one blood RNA sample per participant (by randomly sampling).

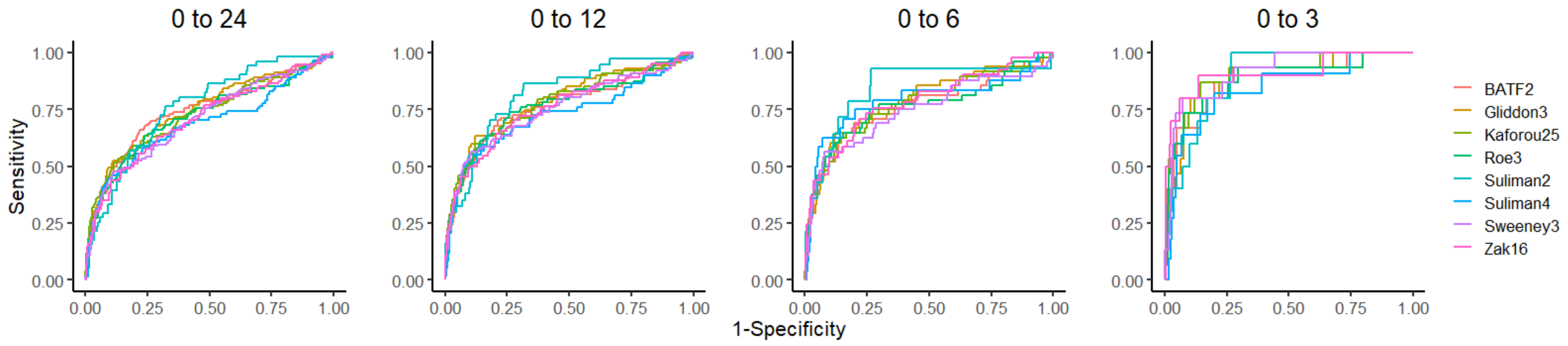

**Supplementary Figure 9.** Diagnostic accuracy of eight best performing transcriptional signatures for incipient tuberculosis (TB) shown in receiver operating characteristic space, stratified by months to disease. Grey shaded zones indicate 95% confidence intervals for each signature. Sensitivity analysis presented using biomarkers cut-offs defined by the maximal Youden indices for each time period, benchmarked against minimal (grey dashed box) and optimal (black dashed box) criteria from the WHO Target Product Profile for incipient TB biomarkers.

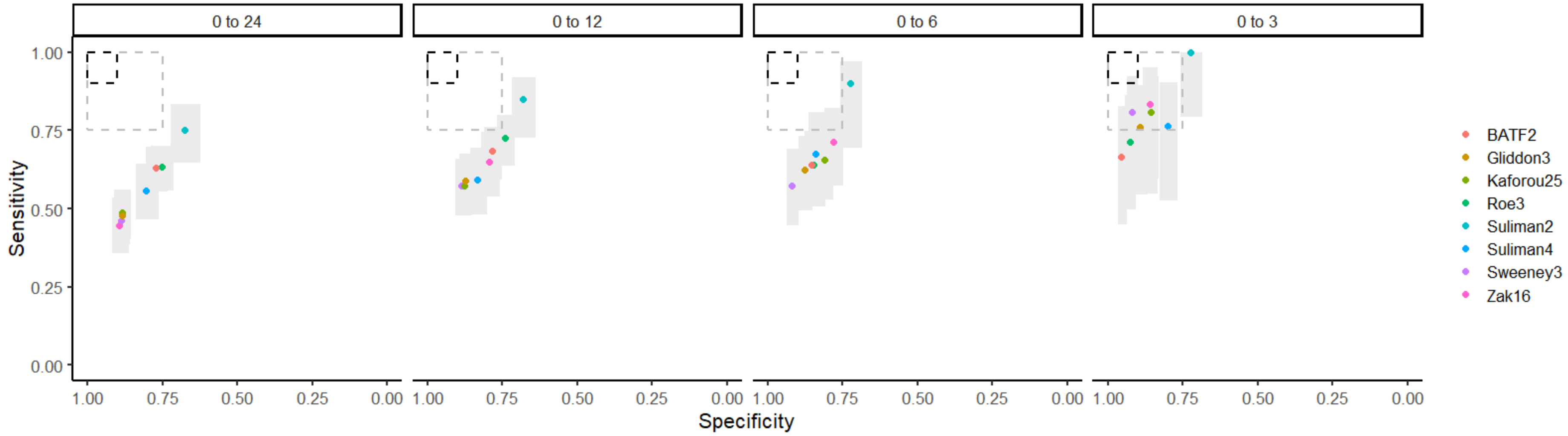

**Supplementary Figure 10.** Receiver operating characteristic curves showing diagnostic accuracy of eight best performing transcriptional signatures for incipient tuberculosis (TB), stratified by months from sample collection to disease. Sensitivity analysis using mutually exclusive time periods of 0-3, 3-6, 6-12 and 12-24 months.

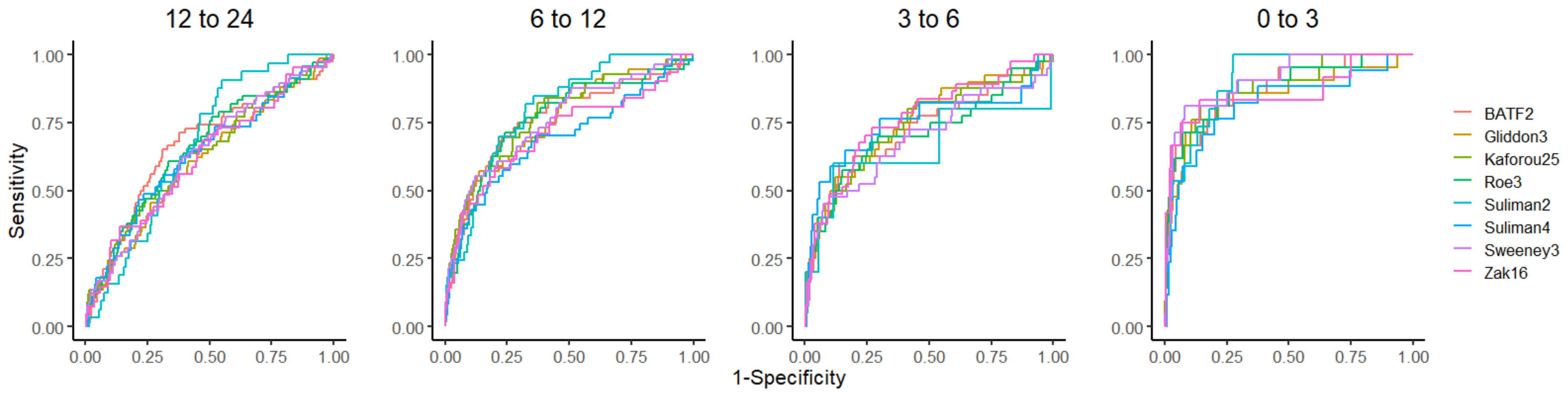

**Supplementary Table 1.** Baseline characteristics of participants in meta-analysis of concise whole blood transcriptional signatures for incipient tuberculosis (TB), stratified by study. ACS = adolescent cohort study; GC6-74 = Bill and Melinda Gates Foundation Grand Challenges 6-74 TB contacts study; IGRA = interferon gamma release assay; IQR = interquartile range. Data shown as n(%) unless otherwise specified.

| Category | Level | London contacts | Leicester contacts | GC6-74 | ACS | Overall |
|---|---|---|---|---|---|---|
| **Patients** | *n* | 324 | 103 | 334 | 144 | 905 |
| **Age** | *Median (IQR)* | 34.00 [26.00, 47.00] | 35.00 [24.00, 45.50] | 23.00 [19.00, 35.00] | 16.00 [15.00, 17.00] | 26.00 [18.00, 40.00] |
| **Gender** | *Female* | 153 (47.2) | 43 (41.7) | 197 (59.0) | 97 (67.4) | 490 (54.1) |
| | *Male* | 166 (51.2) | 60 (58.3) | 137 (41.0) | 47 (32.6) | 410 (45.3) |
| | *Missing* | 5 (1.5) | 0 (0.0) | 0 (0.0) | 0 (0.0) | 5 (0.6) |
| **Samples per patient** | *Median (IQR)* | 1.00 [1.00, 1.00] | 1.00 [1.00, 1.00] | 1.00 [1.00, 1.00] | 1.00 [1.00, 3.00] | 1.00 [1.00, 1.00] |
| **IGRA** | *Negative* | 219 (67.6) | 50 (48.5) | 0 (0.0) | 3 (2.1) | 272 (30.1) |
| | *Positive* | 105 (32.4) | 53 (51.5) | 0 (0.0) | 36 (25.0) | 194 (21.4) |
| | *Missing* | 0 (0.0) | 0 (0.0) | 334 (100.0) | 105 (72.9) | 439 (48.5) |
| **Country** | *Ethiopia* | 0 (0.0) | 0 (0.0) | 36 (10.8) | 0 (0.0) | 36 (4.0) |
| | *South Africa* | 0 (0.0) | 0 (0.0) | 180 (53.9) | 144 (100.0) | 324 (35.8) |
| | *The Gambia* | 0 (0.0) | 0 (0.0) | 118 (35.3) | 0 (0.0) | 118 (13.0) |
| | *UK* | 324 (100.0) | 103 (100.0) | 0 (0.0) | 0 (0.0) | 427 (47.2) |
| **Outcome** | *Non-progressor* | 316 (97.5) | 99 (96.1) | 259 (77.5) | 104 (72.2) | 778 (86.0) |
| | *Incipient TB* | 8 (2.5) | 4 (3.9) | 75 (22.5) | 40 (27.8) | 127 (14.0) |
| **Months from recruitment to TB** | *Median (IQR)* | 8.63 [6.39, 11.08] | 1.92 [1.00, 3.16] | 10.50 [5.50, 17.50] | 14.37 [8.80, 18.62] | 10.27 [5.50, 17.50] |
| **Microbiological confirmation** | *No* | 5 (62.5) | 0 (0.0) | 5 (6.7) | 0 (0.0) | 10 (7.9) |
| | *Yes* | 3 (37.5) | 4 (100.0) | 70 (93.3) | 40 (100.0) | 117 (92.1) |
| **Pulmonary** | *No* | 7 (87.5) | 1 (25.0) | 0 (0.0) | 0 (0.0) | 8 (6.3) |
| | *Yes* | 1 (12.5) | 3 (75.0) | 0 (0.0) | 40 (100.0) | 44 (34.6) |
| | *Missing* | 0 (0.0) | 0 (0.0) | 75 (100.0) | 0 (0.0) | 75 (59.1) |
| **Total samples** | *n* | 8 | 4 | 98 | 73 | 183 |
| **Months from sample to TB** | *Median (IQR)* | 8.63 [6.39, 11.08] | 1.92 [1.00, 3.16] | 7.50 [5.50, 15.50] | 9.33 [6.57, 14.97] | 8.50 [5.50, 15.10] |
| | *<3* | 1 (12.5) | 3 (75.0) | 6 (6.1) | 11 (15.1) | 21 (11.5) |
| | *3 to 6* | 1 (12.5) | 1 (25.0) | 35 (35.7) | 3 (4.1) | 40 (21.9) |
| | *6 to 12* | 5 (62.5) | 0 (0.0) | 23 (23.5) | 28 (38.4) | 56 (30.6) |
| | *>12* | 1 (12.5) | 0 (0.0) | 34 (34.7) | 31 (42.5) | 66 (36.1) |

**Supplementary Table 2.** Validation of reconstructed signature models against the authors' original descriptions by comparing AUCs in common datasets. No validation was possible for the Huang11 model as no AUC reported by authors in their original training / test set

| Signature | Original AUC | Our AUC | Common dataset |
|---|---|---|---|
| Zak16* | 0.69 | 0.70 | Zak test |
| Suliman4[$] | 0.67 | 0.66 | Suliman test |
| Walter45* | 0.98 | 0.98 | Walter test |
| Maertzdorf4[%] | 0.98 | 1.00 | Maertzdorf training |

*Support vector machine models including individual genes with linear kernels using the R kernlab package.

[$]Also reconstructed as a support vector machine model using gene pairs. Since performance was marginally better and closer to the authors' AUC in their own test set using ((GAS6 + SEPT4) - (CD1C + BLK); AUC=0.66), compared to the gene pairs SVM (AUC=0.65), the simple formula approach was used.

[%]Random forest model created using randomForest package in R.

**Supplementary Table 3.** Tables showing receiver operating characteristic areas under the curve (95% confidence intervals) for 17 transcriptional signatures for identification of incipient TB over a two-year period, stratified by (a) study, and (b) study and time interval to disease. ACS = adolescent cohort study; GC6-74 = Bill and Melinda Gates Foundation Grand Challenges 6-74 TB contacts study.

**(a)**

| Signature | ACS | GC6-74 | London contacts | Leicester contacts |
|---|---|---|---|---|
| **Anderson38** | 0.67 (0.59 - 0.75) | 0.56 (0.49 - 0.64) | 0.51 (0.26 - 0.75) | 0.66 (0.42 - 0.9) |
| **BATF2** | 0.81 (0.74 - 0.88) | 0.68 (0.61 - 0.76) | 0.81 (0.61 - 1) | 0.71 (0.33 - 1) |
| **Gjoen7** | 0.72 (0.64 - 0.8) | 0.64 (0.57 - 0.71) | 0.83 (0.71 - 0.94) | 0.66 (0.28 - 1) |
| **Gliddon3** | 0.72 (0.64 - 0.8) | 0.74 (0.67 - 0.8) | 0.84 (0.62 - 1) | 0.66 (0.21 - 1) |
| **Huang11** | 0.68 (0.6 - 0.76) | 0.69 (0.62 - 0.76) | 0.67 (0.48 - 0.87) | 0.65 (0.3 - 0.99) |
| **Kaforou25** | 0.79 (0.72 - 0.86) | 0.7 (0.63 - 0.77) | 0.84 (0.64 - 1) | 0.66 (0.22 - 1) |
| **Maertzdorf4** | 0.75 (0.68 - 0.83) | 0.63 (0.56 - 0.7) | 0.81 (0.67 - 0.95) | 0.66 (0.23 - 1) |
| **NPC2** | 0.64 (0.55 - 0.72) | 0.69 (0.62 - 0.75) | 0.84 (0.71 - 0.97) | 0.8 (0.64 - 0.96) |
| **Qian17** | 0.7 (0.61 - 0.78) | 0.61 (0.54 - 0.68) | 0.81 (0.68 - 0.93) | 0.71 (0.46 - 0.97) |
| **Rajan5** | 0.7 (0.62 - 0.78) | 0.54 (0.47 - 0.61) | 0.45 (0.17 - 0.72) | 0.53 (0.32 - 0.74) |
| **Roe3** | 0.83 (0.76 - 0.89) | 0.64 (0.56 - 0.71) | 0.79 (0.6 - 0.99) | 0.74 (0.33 - 1) |
| **Singhania20** | 0.69 (0.6 - 0.77) | 0.66 (0.59 - 0.73) | 0.73 (0.61 - 0.85) | 0.45 (0.12 - 0.78) |
| **Suliman2** | 0.8 (0.74 - 0.87) | NA | 0.8 (0.63 - 0.98) | 0.62 (0.21 - 1) |
| **Suliman4** | 0.75 (0.67 - 0.83) | 0.63 (0.52 - 0.74) | 0.85 (0.72 - 0.98) | 0.7 (0.28 - 1) |
| **Sweeney3** | 0.78 (0.71 - 0.85) | 0.69 (0.62 - 0.76) | 0.75 (0.55 - 0.95) | 0.65 (0.22 - 1) |
| **Walter45** | 0.59 (0.5 - 0.68) | 0.55 (0.47 - 0.62) | 0.49 (0.27 - 0.7) | 0.52 (0.35 - 0.7) |
| **Zak16** | 0.68 (0.49 - 0.87) | 0.67 (0.6 - 0.74) | 0.79 (0.59 - 0.99) | 0.76 (0.43 - 1) |

**(b)**

| Months to TB | 12 to 24 | | 6 to 12 | | 3 to 6 | | 0 to 3 | |
|---|---|---|---|---|---|---|---|---|
| **Study** | **GC6-74** | **ACS** | **GC6-74** | **ACS** | **GC6-74** | **ACS** | **GC6-74** | **ACS** |
| **Anderson38** | 0.52 (0.42 - 0.62) | 0.55 (0.44 - 0.66) | 0.67 (0.54 - 0.8) | 0.64 (0.53 - 0.75) | 0.59 (0.47 - 0.7) | 0.54 (0.01 - 1) | 0.62 (0.36 - 0.89) | 0.69 (0.55 - 0.84) |
| **BATF2** | 0.64 (0.53 - 0.75) | 0.68 (0.58 - 0.78) | 0.73 (0.6 - 0.87) | 0.79 (0.69 - 0.89) | 0.72 (0.62 - 0.83) | 0.82 (0.57 - 1) | 0.79 (0.55 - 1) | 0.9 (0.82 - 0.98) |
| **Gjoen7** | 0.6 (0.49 - 0.71) | 0.64 (0.54 - 0.74) | 0.7 (0.57 - 0.83) | 0.67 (0.54 - 0.79) | 0.68 (0.58 - 0.77) | 0.76 (0.6 - 0.93) | 0.68 (0.37 - 1) | 0.74 (0.55 - 0.94) |
| **Gliddon3** | 0.64 (0.52 - 0.75) | 0.56 (0.46 - 0.66) | 0.8 (0.68 - 0.92) | 0.72 (0.63 - 0.81) | 0.76 (0.67 - 0.85) | 0.76 (0.45 - 1) | 0.81 (0.57 - 1) | 0.85 (0.7 - 1) |
| **Huang11** | 0.54 (0.42 - 0.66) | 0.52 (0.42 - 0.62) | 0.82 (0.72 - 0.91) | 0.68 (0.59 - 0.78) | 0.62 (0.51 - 0.73) | 0.72 (0.28 - 1) | 0.88 (0.76 - 1) | 0.71 (0.55 - 0.88) |
| **Kaforou25** | 0.62 (0.51 - 0.74) | 0.63 (0.53 - 0.73) | 0.78 (0.66 - 0.91) | 0.77 (0.7 - 0.85) | 0.73 (0.64 - 0.83) | 0.81 (0.57 - 1) | 0.84 (0.63 - 1) | 0.89 (0.78 - 1) |
| **Maertzdorf4** | 0.57 (0.46 - 0.68) | 0.63 (0.53 - 0.73) | 0.76 (0.64 - 0.87) | 0.73 (0.62 - 0.85) | 0.65 (0.54 - 0.75) | 0.86 (0.73 - 0.98) | 0.71 (0.4 - 1) | 0.85 (0.73 - 0.97) |
| **NPC2** | 0.61 (0.51 - 0.71) | 0.59 (0.47 - 0.71) | 0.68 (0.53 - 0.83) | 0.63 (0.52 - 0.74) | 0.73 (0.64 - 0.81) | 0.67 (0.32 - 1) | 0.9 (0.8 - 1) | 0.69 (0.5 - 0.89) |
| **Qian17** | 0.57 (0.45 - 0.69) | 0.56 (0.45 - 0.68) | 0.7 (0.59 - 0.82) | 0.73 (0.62 - 0.84) | 0.63 (0.52 - 0.74) | 0.79 (0.48 - 1) | 0.78 (0.57 - 0.99) | 0.77 (0.63 - 0.91) |
| **Rajan5** | 0.5 (0.39 - 0.6) | 0.54 (0.44 - 0.65) | 0.47 (0.34 - 0.6) | 0.72 (0.61 - 0.82) | 0.52 (0.41 - 0.62) | 0.68 (0.31 - 1) | 0.64 (0.35 - 0.93) | 0.77 (0.61 - 0.93) |
| **Roe3** | 0.58 (0.47 - 0.7) | 0.72 (0.64 - 0.81) | 0.7 (0.56 - 0.84) | 0.82 (0.74 - 0.89) | 0.71 (0.6 - 0.81) | 0.82 (0.53 - 1) | 0.77 (0.51 - 1) | 0.91 (0.85 - 0.98) |
| **Singhania20** | 0.64 (0.54 - 0.75) | 0.62 (0.5 - 0.74) | 0.65 (0.52 - 0.78) | 0.63 (0.51 - 0.75) | 0.7 (0.62 - 0.78) | 0.65 (0.44 - 0.86) | 0.65 (0.38 - 0.93) | 0.82 (0.65 - 0.98) |
| **Sweeney3** | 0.59 (0.48 - 0.7) | 0.62 (0.53 - 0.71) | 0.78 (0.66 - 0.89) | 0.71 (0.61 - 0.82) | 0.69 (0.58 - 0.8) | 0.81 (0.5 - 1) | 0.91 (0.77 - 1) | 0.9 (0.79 - 1) |
| **Walter45** | 0.5 (0.4 - 0.6) | 0.5 (0.38 - 0.61) | 0.54 (0.41 - 0.66) | 0.55 (0.43 - 0.67) | 0.69 (0.59 - 0.79) | 0.55 (0.31 - 0.8) | 0.52 (0.19 - 0.86) | 0.47 (0.3 - 0.63) |

**Supplementary Table 4.** Receiver operating characteristic areas under the curve (95% confidence intervals) showing diagnostic accuracy of candidate transcriptional signatures for incipient TB, stratified by time interval to disease, among pooled dataset of four contributing studies. P values represent paired comparisons against the best performing signature available for all participants (BATF2) over two years.

| Signature | 0 to 24 | p | 0 to 12 | 0 to 6 | 0 to 3 |
|---|---|---|---|---|---|
| **Suliman2** | 0.77 (0.71 - 0.82) | 0.355 | 0.82 (0.76 - 0.88) | 0.85 (0.74 - 0.95) | 0.91 (0.86 - 0.96) |
| **BATF2** | 0.74 (0.69 - 0.78) | ref | 0.77 (0.72 - 0.82) | 0.78 (0.7 - 0.85) | 0.87 (0.79 - 0.95) |
| **Kaforou25** | 0.73 (0.69 - 0.78) | 0.852 | 0.78 (0.73 - 0.83) | 0.79 (0.72 - 0.86) | 0.88 (0.8 - 0.97) |
| **Gliddon3** | 0.73 (0.68 - 0.77) | 0.575 | 0.77 (0.72 - 0.82) | 0.78 (0.71 - 0.85) | 0.85 (0.74 - 0.96) |
| **Sweeney3** | 0.72 (0.68 - 0.77) | 0.438 | 0.77 (0.71 - 0.82) | 0.77 (0.69 - 0.84) | 0.91 (0.84 - 0.97) |
| **Roe3** | 0.72 (0.67 - 0.77) | 0.109 | 0.77 (0.71 - 0.82) | 0.77 (0.7 - 0.84) | 0.88 (0.79 - 0.97) |
| **Suliman4** | 0.7 (0.64 - 0.76) | 0.257 | 0.73 (0.66 - 0.8) | 0.78 (0.68 - 0.89) | 0.82 (0.69 - 0.94) |
| **Zak16** | 0.7 (0.64 - 0.76) | 0.939 | 0.76 (0.69 - 0.82) | 0.79 (0.71 - 0.86) | 0.86 (0.71 - 1) |
| **NPC2** | 0.68 (0.64 - 0.73) | 0.012 | 0.71 (0.66 - 0.77) | 0.75 (0.69 - 0.82) | 0.78 (0.66 - 0.9) |
| **Maertzdorf4** | 0.68 (0.63 - 0.73) | 0.001 | 0.73 (0.68 - 0.78) | 0.71 (0.64 - 0.79) | 0.8 (0.69 - 0.91) |
| **Gjoen7** | 0.67 (0.63 - 0.72) | 0.001 | 0.69 (0.64 - 0.75) | 0.7 (0.62 - 0.77) | 0.75 (0.61 - 0.88) |
| **Singhania20** | 0.67 (0.62 - 0.72) | 0.006 | 0.68 (0.62 - 0.73) | 0.72 (0.65 - 0.78) | 0.74 (0.6 - 0.87) |
| **Huang11** | 0.67 (0.62 - 0.71) | 0.013 | 0.71 (0.66 - 0.76) | 0.68 (0.6 - 0.75) | 0.75 (0.64 - 0.86) |
| **Qian17** | 0.66 (0.61 - 0.71) | <0.0001 | 0.71 (0.66 - 0.76) | 0.7 (0.63 - 0.77) | 0.79 (0.7 - 0.88) |
| **Anderson38** | 0.6 (0.55 - 0.65) | <0.0001 | 0.64 (0.58 - 0.7) | 0.64 (0.56 - 0.72) | 0.71 (0.6 - 0.82) |
| **Rajan5** | 0.55 (0.5 - 0.6) | <0.0001 | 0.59 (0.53 - 0.65) | 0.57 (0.49 - 0.66) | 0.68 (0.56 - 0.81) |
| **Walter45** | 0.55 (0.5 - 0.6) | <0.0001 | 0.58 (0.53 - 0.64) | 0.62 (0.54 - 0.69) | 0.47 (0.34 - 0.6) |

**Supplementary Table 5.** Diagnostic accuracy of eight best performing transcriptional signatures for incipient tuberculosis (TB), stratified by months to disease. Positive and negative predictive values (PPVs/NPVs) shown assuming 2% pre-test probability. Data presented as estimate (95% confidence interval). Data presented graphically in Figure 4.

| **Signature** | **Sensitivity** | **Specificity** | **PPV** | **NPV** |
|---|---|---|---|---|
| **Timeframe** | **0 to 24** | | | |
| **BATF2** | 0.35 (0.29 - 0.43) | 0.93 (0.91 - 0.95) | 0.074 (0.049 - 0.108) | 0.986 (0.984 - 0.988) |
| **Gliddon3** | 0.31 (0.25 - 0.39) | 0.95 (0.93 - 0.97) | 0.085 (0.054 - 0.128) | 0.986 (0.984 - 0.987) |
| **Kaforou25** | 0.35 (0.28 - 0.42) | 0.95 (0.93 - 0.97) | 0.094 (0.06 - 0.138) | 0.986 (0.985 - 0.988) |
| **Roe3** | 0.3 (0.24 - 0.37) | 0.94 (0.92 - 0.96) | 0.07 (0.044 - 0.105) | 0.985 (0.983 - 0.987) |
| **Suliman2** | 0.25 (0.17 - 0.35) | 0.95 (0.92 - 0.97) | 0.068 (0.034 - 0.123) | 0.984 (0.982 - 0.987) |
| **Suliman4** | 0.29 (0.21 - 0.37) | 0.95 (0.93 - 0.97) | 0.08 (0.046 - 0.129) | 0.985 (0.983 - 0.987) |
| **Sweeney3** | 0.4 (0.33 - 0.47) | 0.92 (0.9 - 0.94) | 0.077 (0.052 - 0.11) | 0.987 (0.985 - 0.989) |
| **Zak16** | 0.33 (0.25 - 0.42) | 0.94 (0.91 - 0.95) | 0.074 (0.045 - 0.115) | 0.986 (0.983 - 0.988) |
| **Timeframe** | **0 to 12** | | | |
| **BATF2** | 0.43 (0.34 - 0.52) | 0.92 (0.9 - 0.94) | 0.083 (0.056 - 0.116) | 0.988 (0.985 - 0.99) |
| **Gliddon3** | 0.41 (0.33 - 0.5) | 0.94 (0.92 - 0.95) | 0.093 (0.062 - 0.132) | 0.987 (0.985 - 0.989) |
| **Kaforou25** | 0.44 (0.36 - 0.53) | 0.94 (0.92 - 0.96) | 0.103 (0.07 - 0.143) | 0.988 (0.986 - 0.99) |
| **Roe3** | 0.37 (0.29 - 0.46) | 0.94 (0.92 - 0.95) | 0.085 (0.056 - 0.123) | 0.986 (0.984 - 0.989) |
| **Suliman2** | 0.34 (0.23 - 0.47) | 0.95 (0.92 - 0.96) | 0.084 (0.046 - 0.14) | 0.986 (0.983 - 0.989) |
| **Suliman4** | 0.33 (0.24 - 0.44) | 0.95 (0.93 - 0.96) | 0.089 (0.052 - 0.138) | 0.986 (0.984 - 0.988) |
| **Sweeney3** | 0.52 (0.43 - 0.61) | 0.92 (0.89 - 0.93) | 0.092 (0.065 - 0.124) | 0.989 (0.987 - 0.992) |
| **Zak16** | 0.42 (0.32 - 0.53) | 0.94 (0.92 - 0.95) | 0.096 (0.061 - 0.142) | 0.988 (0.985 - 0.99) |
| **Timeframe** | **0 to 6** | | | |
| **BATF2** | 0.51 (0.39 - 0.63) | 0.93 (0.91 - 0.94) | 0.099 (0.065 - 0.139) | 0.989 (0.986 - 0.992) |
| **Gliddon3** | 0.43 (0.31 - 0.55) | 0.94 (0.92 - 0.95) | 0.093 (0.058 - 0.137) | 0.988 (0.985 - 0.99) |
| **Kaforou25** | 0.48 (0.36 - 0.6) | 0.94 (0.92 - 0.95) | 0.102 (0.066 - 0.147) | 0.989 (0.986 - 0.991) |
| **Roe3** | 0.44 (0.33 - 0.57) | 0.94 (0.92 - 0.95) | 0.098 (0.062 - 0.143) | 0.988 (0.985 - 0.991) |
| **Suliman2** | 0.5 (0.3 - 0.7) | 0.94 (0.92 - 0.95) | 0.11 (0.055 - 0.178) | 0.989 (0.985 - 0.994) |
| **Suliman4** | 0.47 (0.31 - 0.63) | 0.95 (0.93 - 0.96) | 0.117 (0.066 - 0.18) | 0.989 (0.985 - 0.992) |
| **Sweeney3** | 0.57 (0.45 - 0.69) | 0.91 (0.89 - 0.93) | 0.095 (0.065 - 0.129) | 0.991 (0.988 - 0.993) |
| **Zak16** | 0.45 (0.32 - 0.59) | 0.94 (0.92 - 0.95) | 0.102 (0.061 - 0.153) | 0.988 (0.985 - 0.991) |
| **Timeframe** | **0 to 3** | | | |
| **BATF2** | 0.67 (0.45 - 0.83) | 0.93 (0.91 - 0.94) | 0.126 (0.076 - 0.175) | 0.993 (0.988 - 0.996) |
| **Gliddon3** | 0.52 (0.32 - 0.72) | 0.94 (0.92 - 0.95) | 0.112 (0.06 - 0.171) | 0.99 (0.985 - 0.994) |
| **Kaforou25** | 0.62 (0.41 - 0.79) | 0.94 (0.92 - 0.95) | 0.129 (0.075 - 0.185) | 0.992 (0.987 - 0.996) |
| **Roe3** | 0.62 (0.41 - 0.79) | 0.94 (0.92 - 0.95) | 0.132 (0.077 - 0.189) | 0.992 (0.987 - 0.996) |
| **Suliman2** | 0.53 (0.3 - 0.75) | 0.94 (0.92 - 0.95) | 0.117 (0.056 - 0.188) | 0.99 (0.985 - 0.995) |
| **Suliman4** | 0.47 (0.26 - 0.69) | 0.95 (0.93 - 0.96) | 0.117 (0.056 - 0.193) | 0.989 (0.984 - 0.993) |
| **Sweeney3** | 0.81 (0.6 - 0.92) | 0.91 (0.89 - 0.93) | 0.129 (0.085 - 0.166) | 0.996 (0.991 - 0.998) |
| **Zak16** | 0.67 (0.39 - 0.86) | 0.94 (0.92 - 0.95) | 0.144 (0.074 - 0.21) | 0.993 (0.987 - 0.997) |
